# Gal1 repression memory in budding yeast

**DOI:** 10.1101/2022.02.03.478948

**Authors:** Lea Schuh, Igor Kukhtevich, Poonam Bheda, Melanie Schulz, Maria Bordukova, Robert Schneider, Carsten Marr

## Abstract

Cells must continuously adapt to changing environments and, thus, have evolved mechanisms allowing them to respond to repeated stimuli. For example, faster gene induction upon a repeated stimulus aids adaptation - a process known as reinduction memory. However, whether such a memory exists for gene repression is unclear. Here, we studied gene repression across repeated carbon source shifts in over 2,500 single *Saccharomyces cerevisiae* cells. By monitoring the expression of a carbon source-responsive gene, galactokinase 1 (*Gal1*), and mathematical modeling, we discovered repression memory at the population and single-cell level. Using a repressor model to estimate single-cell repression parameters, we show that repression memory is due to a shortened repression delay, the estimated time gap between carbon source shift and Gal1 expression termination, upon the repeated carbon source shift. Additionally, we show that cells lacking *Elp6* display a gain-of-repression-memory phenotype characterized by a stronger decrease in repression delay between two consecutive carbon source shifts. Collectively, our study provides the first quantitative description of repression memory in single cells.

## INTRODUCTION

Cells receive and process external signals to optimally adapt to changing environments. Repeated stimulation from the same external signal induces an adapted transcriptional response, a phenomenon termed transcriptional memory (*1*). It is crucial to understand the mechanisms underlying transcriptional memory due to its implications for a broad range of cellular functions, including the human adaptive immune system (*2, 3*), disease development in diabetes (*4, 5*), and aging (*6*). However, transcriptional memory has primarily been researched concerning gene induction despite gene repression playing an essential role in gene regulation (*7, 8*). This raises the question of whether memory exists in repression.

The adaptation of *Saccharomyces cerevisiae* (budding yeast) to carbon sources is among the most well-studied eukaryotic signal integration systems. Whereas glucose directly enters glycolysis, a vital metabolic route providing cells with energy, galactose is first converted to glucose-6-phosphate (*9, 10*), necessitating the production of Gal gene-encoded enzymes (*11*). Repeated alternations between glucose and galactose media revealed that yeast cells are primed by their carbon source history, exhibiting transcriptional memory: repeated galactose induction results in enhanced Gal gene expression (*12–16*). Bheda et al. examined the expression of galactokinase 1 (*Gal1*) in single cells, for which reinduction memory has been well characterized, and discovered that a shorter delay, rather than an increased expression rate, contributed to the observed increase in Gal1 levels (*17*). Moreover, they identified *elp6Δ* as a gain-of-reinduction memory mutant, with *elp6Δ* cells showing Gal1 levels comparable to wildtype cells in the first induction, but earlier induction onset and increased Gal1 levels in the second induction. While, as these examples show, reinduction memory upon galactose induction has been thoroughly researched, and, while, Lee et al. suggested repression memory for bulk populations (*18*), it is uncertain if individual cells display Gal1 repression memory upon repeated glucose repression (Figure 1).

**Figure 1.**
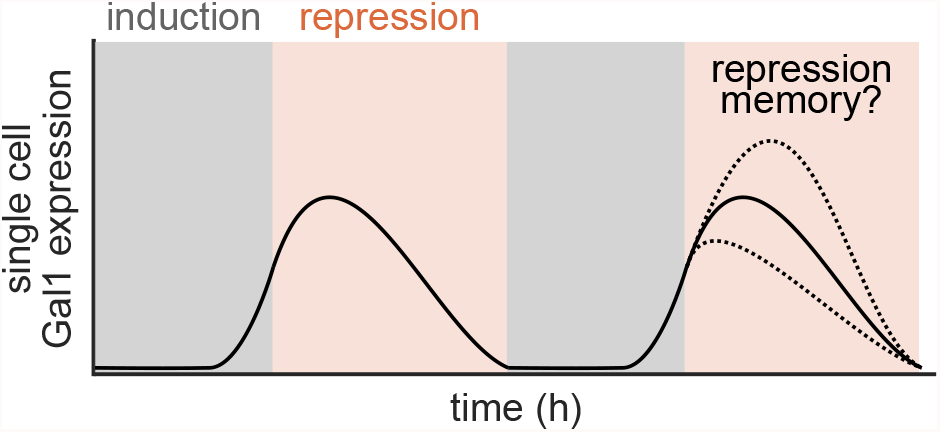
Does gene repression memory exist in budding yeast? Previous studies focused on gene expression during induction. However, whether there is memory in repression indicated by altered repression kinetics is still unknown.

We measured Gal1 expression in wildtype budding yeast cells via a Gal1-GFP (green fluorescent protein) fusion across repeated galactose inductions and glucose repressions using time-lapse microscopy coupled with a microfluidic device to follow and study Gal1 repression kinetics. In the second repression, we discovered that the time of maximal mean Gal1 expression was shortened. By using a mathematical model to quantify single-cell Gal1-GFP kinetics during glucose repression, and by distinguishing between repressor and non-repressor cells, we revealed that the shortened time to maximal mean Gal1 expression at the population level was not caused by different fractions of repressor cells between consecutive repressions. Using the estimated single-cell parameters, we found the repression delay, which is the estimated time gap between the galactose to glucose shift and the Gal1 expression termination, to be shortened in the second repression at the single-cell level, implying Gal1 repression memory. Furthermore, we repeated the experiments and analysis for the gain-of-induction memory mutant *elp6Δ*. Remarkably, *elp6Δ* cells showed a stronger repression memory effect than wildtype, making *elp6Δ* also a gain-of-repression memory mutant.

## RESULTS

### Automated time-lapse microscopy and microfluidics allow for the quantification of single-cell Gal1 repression kinetics across repeated carbon source shifts

To study Gal1 repression kinetics over multiple repressions, we exposed wildtype budding yeast cells alternatingly to glucose or galactose media (Figure 2A). For this, we cultured the cells in custom-made microfluidic devices to ensure precise media shifts and long-term tracking (*17*). Gal1 expression levels in single yeast cells were monitored using a Gal1-GFP fusion, a standard reporter to study gene expression in time-lapse microscopy (see Bheda et al., 2020 for details). We captured images from the microfluidics chambers every 3 min totaling 320 images per chamber during a 16-h experiment. The yeast cells were then semi-automatically segmented, mapped and the total Gal1-GFP fluorescence signal per cell and time point was extracted using Autotrack and PhyloCell (*19*), YeaZ (*20*) and Cell-ACDC (*21*) (see Materials and Methods), yielding over 2,500 single-cell Gal1 expression traces (Figure 2B). Asymmetric budding allowed us to identify mother-daughter relationships. As expected, the total Gal1-GFP fluorescence signal of single cells reveals Gal1 inductions and repressions during galactose and glucose, respectively, and increased overall Gal1 levels in induction i2 (Figure 2B).

**Figure 2.**
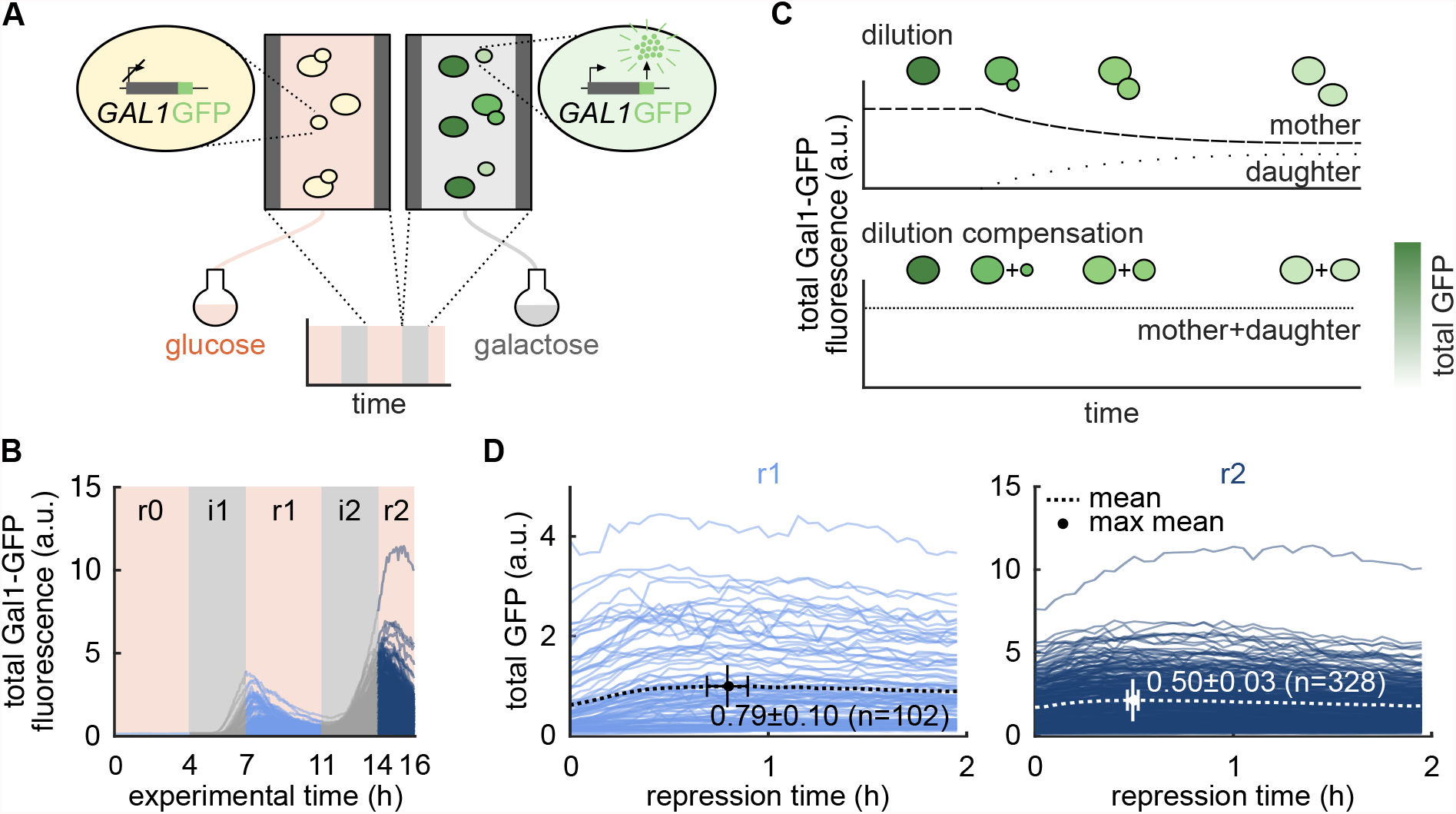
A shortened time to maximal mean total GFP in the second repression at the population level. (A) Budding yeast cells were grown in microfluidic chambers and alternatingly exposed to a medium containing either glucose (orange) or galactose (gray) as carbon source. Galactokinase 1 (*Gal1*) is induced in cells exposed to galactose and repressed in cells exposed to glucose. Gal1 expression was monitored via a Gal1-GFP fusion and time-lapse microscopy. (B) Single-cell traces of total Gal1-GFP fluorescence signal across two inductions i1 and i2 (gray) and repressions r0, r1, and r2 (blue). (C) Budding decreases the total Gal1-GFP fluorescence signal in mother cells and increases the total Gal1-GFP fluorescence signal in daughter cells (top). To compensate for this dilution, we summed up the total Gal1-GFP fluorescence signal of each mother cell present and its progeny during one repression (bottom). (D) Single-cell traces of total GFP signal adjusted for dilution (see (C)) for the first two hours of repressions r1 (left) and r2 (right). Time to maximal mean total GFP is 17 min shorter in repression r2, where mean total GFP is indicated by the dotted line and the maximal mean total GFP is highlighted by the dot. Bootstrap (10^5^) samples were drawn to generate mean ± std. The replicate analysis can be found in Figure S1.

During budding, cytoplasmic proteins are disseminated between the mother and daughter cells. Assuming a constant Gal1 protein amount, its redistribution decreases the total Gal1-GFP fluorescence signal in the mother cell (Figure 2C top), a phenomenon called dilution. To deconvolute dilution and repression kinetics, we calculated the sum of the total Gal1-GFP fluorescence signal of the mother cell and its progeny (Figure 2C bottom, and see Materials and Methods). In the following, the adjusted sum of the total Gal1-GFP fluorescence signal of the mother cell and its progeny is referred to as total GFP. We applied the same dilution compensation to repressions r1 and r2 (Figure 2D).

### Decreased time to maximal total GFP in repression r2 at the population level

Following a galactose-glucose shift, total GFP intensities initially rise before decreasing (Figure 2D). To determine the repression kinetics at the population level, we calculated the mean total GFP signal over time for repressions r1 and r2. Interestingly, the time to attain the maximal mean total GFP reduced from 0.79 ± 0.10 h (mean ± std, n = 102 cells) in r1 to 0.50 ± 0.03 h (n = 328 cells) in r2, where the time point of the maximal mean was bootstrapped 10^5^ times (Figure 2D). This demonstrates a decreased time to maximal total GFP in r2 at the population level.

### The computational model distinguishes between repressor and non-repressor cells

At the single-cell level, Gal1 induction delay varies significantly. Zacharioudakis et al. showed that Gal1 induction caused by a glucose–galactose media shift results in a bimodal population distribution, with only a subset of cells inducing Gal1 even after several hours of galactose exposure (*16*). As in our experiments, repression was preceded by 3 h of galactose induction, we expected that our cell population at the start of repression contained induced and uninduced cells, which show and do not show repression kinetics, respectively. However, independent of repression memory, a larger proportion of non-repressor cells in repression r1 could explain the decrease in the time to maximal mean total GFP in repression r2 (Figure 2D). As it is difficult to distinguish between repressor and non-repressor cells from the total GFP traces alone (Figure 2D), we used computational modeling and model selection to systematically describe the kinetics of single total GFP traces. Since Gal1 induction results in an approximate 1000-fold change in Gal1 expression (*22*), we assumed that stochasticity inherent to gene expression was insignificant and that a deterministic modeling approach was sufficient in describing the kinetics of the total GFP traces. To discriminate between repressor and non-repressor cells, we defined two models. The non-repressor model assumes a constant basal GFP production and degradation over time with rates r_basal_and r_deg_ (Figure 3A left), since we observed a gradual increase in total GFP signal in cells visually identified as not showing repression kinetics (Figure 3B top right). The temporal variation of the total GFP signal over time indicated by the non-repressor model is summarized using the following ordinary differential equation:

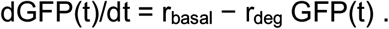

**Figure 3.**
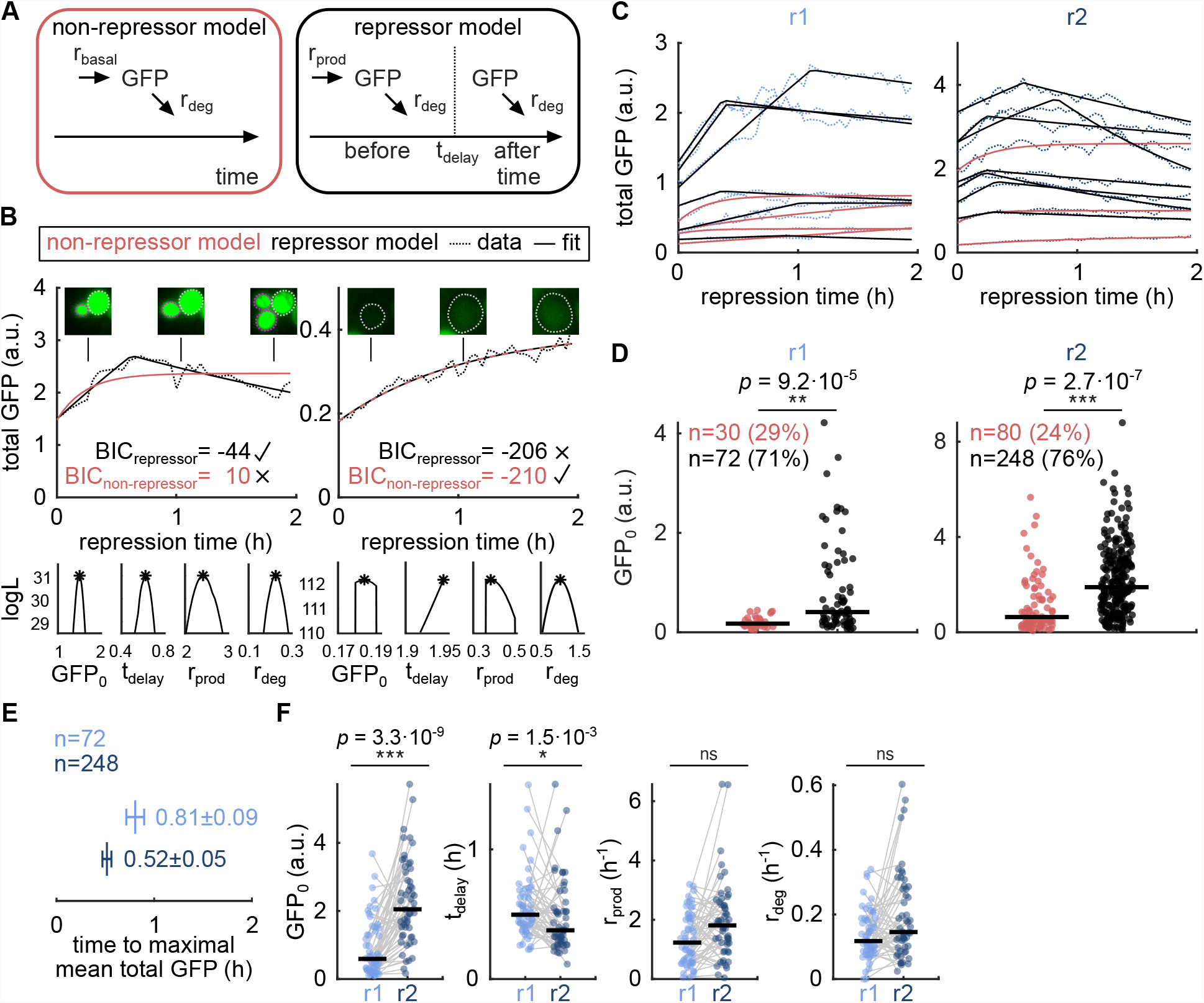
Shortened repression delay in repression r2 at the single-cell level. (A) Left: a model for non-repressor cells composed of basal GFP production (r_basal_) and degradation (r_deg_). Right: a model for repressor cells composed of an initial constant and active GFP production (r_prod_) and degradation (r_deg_) until a delayed repression onset (t_delay_) where GFP production is switched off. (B) Top: two exemplary total GFP traces (dotted line) and fits of the non-repressor model (red solid line) and repressor model (black solid line). Exemplary images of the cell(s) at three different time points are shown above. Mother cells are circled in gray, progeny in pink. The better fitting model was selected according to the Bayesian information criterion (BIC). Left: total GFP trace better fitted by the repressor model. Right: total GFP trace fitted equally well by the non-repressor and repressor model. Due to the higher model complexity of the repressor model, the repressor model is still rejected. Bottom: profile likelihoods of the repressor model corresponding to the two exemplary total GFP traces above endorse parameter identifiability. Asterisks represent optimized parameters and corresponding log-likelihood (logL) values. (C) Ten exemplary total GFP traces (dotted lines) and best fits (solid lines) for repressions r1 (left) and r2 (right). GFP traces best fitted with a repressor model are shown in black and fits of total GFP traces best fitted with a non-repressor model are shown in red. (D) The median initial total GFP, GFP_0_, is significantly higher (*p* = 9.2·10^−5^ and *p* = 2.7·10^−7^, two-sided median test corrected for multiple testing with Bonferroni correction, m = 20) in traces better fitted by the repressor model (black) than in traces better fitted by the non-repressor model (red). This confirms that the repressor model fits induced cells better, while the non-repressor model fits uninduced cells. The number of cells and percentages of all GFP traces best fitted by the repressor model and non-repressor model are shown. (E) Time to maximal mean total GFP is decreased in repression r2 for repressor cells (0.81 ± 0.09 vs. 0.52 ± 0.05). Bootstrap (10^5^) samples of the repressor cells were drawn to generate mean ± std. (F) Comparison of paired estimated single-cell parameters of repression r1 and r2 shows that the median initial total GFP, GFP_0_, and median repression delay, t_delay_, are significantly different (*p* = 3.3·10^−9^ and *p* = 1.5·10^−3^, respectively, two-sided paired sign test correcting for multiple testing with Bonferroni correction, m = 20, and the number of paired cells = 54), with median GFP_0_ increased and median t_delay_ decreased (median t_delay_ values of 0.50 and 0.38 for r1 and r2, respectively). Median production rate, r_prod_, and median degradation rate, r_deg_, are not significantly different between r1 and r2 (*p* = 0.22 and *p* = 0.34, respectively). The replicate analysis can be found in Figure S1.

This is solved by

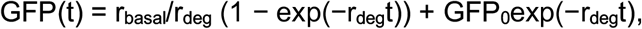

where GFP_0_ = GFP(0), the initial total GFP at time point 0. According to the repressor model, cells that induced Gal1 during galactose induction required time to reestablish glucose-mediated repression. Hence, GFP is actively generated at rate r_prod_ till a time point t_delay_. GFP production is switched off (r_prod_ = 0) and GFP is degraded with rate r_deg_ after this estimated repression delay t_delay_ (Figure 3A right). An example of a cell visually identified as showing repression kinetics can be found in Figure 3B top left. Until t_delay_, the repressor model equals the non-repressor model. The temporal change of total GFP over time described by the repressor model is summarized by the following ordinary differential equations:

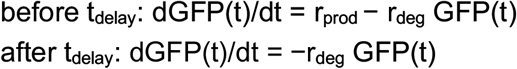

with solutions

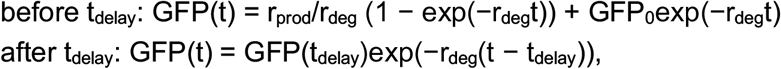

where GFP(t_delay_) = r_prod_/r_deg_(1 -exp (-r_deg_t_delay_)) + GFP_0_ exp (-r_deg_t_delay_). The non-repressor and repressor model comprise four and five model parameters, respectively: initial total GFP, GFP_0_, basal GFP production rate, r_basal_, or GFP production rate, r_prod_, GFP degradation rate r_deg_, a noise parameter σ determining the width of the Gaussian noise distribution (see Materials and Methods), and the repression delay t_delay_ for the repressor model. For repressions r1 and r2, respectively, we performed multi-start maximum likelihood optimization and model selection on both models for each total GFP trace (Figure 3B top and 3C). Calculating the profile likelihoods of exemplary total GFP traces, we found the model parameters of the repressor model to be identifiable (Figure 3B bottom). We then determined whether active repression, i.e. the repressor model, was required to explain a total GFP trace using the Bayesian information criterion (BIC). A BIC difference of ten between the repressor and non-repressor model (BIC_repressor_ < BIC_non-repressor_ – 10) was considered an appropriate threshold to reject the non-repressor model with fewer model parameters (see Materials and Methods and Figure 3B). Of all total GFP traces, 71% and 76% of r1 and r2, respectively, were discovered to require the repressor model (Figure 3D) and are henceforth referred to as “repressor cells.” The higher proportion of repressor cells was expected due to Gal1 transcriptional reinduction memory resulting in increased proportions of cells producing detectable GFP in induction i2. The median initial total GFP, GFP_0_, was significantly higher in repressor cells than in non-repressor cells (*p* = 9.2·10^−2^ for r1 and *p* = 2.7·10^−3^ for r2) (Figure 3D). This implies that our repression models can discriminate between cells that were repressing Gal1 and cells uninduced at the beginning of repression. The overlapping ranges of initial total GFP between repressor and non-repressor cells reveal how simple thresholding could result in wrong differentiation between repressor and non-repressor cells.

### Decreased time to maximal total GFP is also present in the repressor cell subpopulation

To determine whether the previously described decrease in time to maximal mean total GFP in r2 is due to a different fraction of non-repressor cells between r1 and r2, we computed the times to maximal mean total GFP on the repressor cell subpopulation. We again found a shortened time to maximal mean total GFP in r2, with 0.81 ± 0.09 (mean ± std, n = 72 cells) and 0.52 ± 0.05 (n = 248 cells) h for r1 and r2, respectively (Figure 3E), demonstrating that the decrease in time to maximal total GFP in r2 is not due to a different fraction of non-repressor cells.

### Shortened repression delay in repression r2 at the single-cell level

To address if the repression kinetics are different between r1 and r2 in individual cells, we compared the paired estimated single-cell parameters of repressor cells present in both repressions. We discovered that the median initial total GFP, GFP_0_, and median repression delay, t_delay_, are substantially different (*p* = 3.3·10^−4^ and *p* = 1.5·10^−5^, respectively) between both repressions using a two-sided paired sign test and multiple test correction (Figure 3F). Median GFP_0_ is increased (median values of 0.59 and 2.05 for r1 and r2, respectively), while median t_delay_ is shortened in r2 (median values of 0.50 and 0.38 for r1 and r2, respectively, where 72% of paired cells showed a decrease in t_delay_), in line with the previously identified decrease in the time to the maximal mean total GFP in r2 at the repressor subpopulation level (Figure 3E). The increased GFP_0_ in r2 conforms to the transcriptional reinduction memory of Gal1 and results in higher GFP_0_ at the start of r2. The median r_prod_ and median r_deg_ between the two repressions were comparable (*p* = 0.22 and *p* = 0.34, respectively) (Figure 3F). In line with our findings, Bheda et al. also identified similar production rates (*17*). We repeated the entire wildtype analysis based on data from independent experiments (see Methods for details). This replicate analysis confirms the conclusions presented in Figures 2B, 2D and 3, in particular the earlier repression response in repression r2 (Figure S1).

### Earlier repression response in repression r2 for *elp6Δ* cells

Intrigued by the findings in wildtype yeast cells, we repeated our analysis for the previously identified gain-of-reinduction memory mutant *Elp6* (*elp6Δ*). The total Gal1-GFP fluorescence signal of single *elp6Δ* cells demonstrates Gal1 induction and repression during galactose and glucose and reinduction memory (Figure 4A). We performed dilution compensation (Figure 2C) on repressions r1 and r2 (Figure 4B) and calculated the mean total GFP signals over time to determine the repression kinetics at the population level. Similar to wildtype cells, we discovered that the time to attain the maximal mean total GFP decreased from 1.40 ± 0.09 h (n = 66 cells) in r1 to 0.71 ± 0.07 h (n = 237 cells) in r2, (Figure 4B). Subsequently, we repeated multi-start maximum likelihood optimization and model selection for the non-repressor and repressor models to show that the shortened time to maximal mean total GFP in r2 is not caused by a larger proportion of non-repressor cells in r1 (Figure 4C). Of all total GFP traces, 64% and 83% of *elp6Δ* cells in r1 and r2 require the repressor model (Figure 4D). When determining the time to maximal mean total GFP for the *elp6Δ* repressor subpopulation, we again identified a shortened time to maximal mean total GFP, with 1.20 ± 0.14 (n = 42 cells) and 0.69 ± 0.07 (n = 196 cells) h for r1 and r2, respectively (Figure 4E). To investigate the earlier repression response at the single-cell level, we compared the paired estimated single-cell parameters of *elp6Δ* repressor cells present in both repressions. We found the median GFP_0_, and median t_delay_, to be significantly different (*p* = 1.2·10^−10^ and *p* = 3.9·10^−5^) between repressions (Figure 4F). Similar to wildtype cells, the median GFP_0_ increased (median values of 0.57 and 3.69 for r1 and r2, respectively), while the median t_delay_ is shortened in r2 (median values of 0.95 and 0.50 for r1 and r2, respectively, where 85% of paired cells showed a decrease in t_delay_), as previously identified at the repressor subpopulation level. The increased initial total GFP level in r2 was expected due to the previously identified reinduction memory of *elp6Δ* cells. Contrary to wildtype cells, we discovered that for *elp6Δ* cells the median production rate, r_prod_, is significantly different between r1 and r2 (*p* = 6.9·10^−7^) with increased median r_prod_ in r2 (median values of 1.10 and 3.10 for r1 and r2, respectively, Figure 4F) reflecting the gain-of-reinduction-memory phenotype reported by Bheda et al. (*17*). Median r_deg_ was comparable between repressions (*p* = 0.39). We repeated the *elp6Δ* analysis based on additional data from independent experiments (see Methods for details). The replicate analysis confirms the conclusions presented in Figure 4, in particular the earlier repression response in repression r2 (Figure S2).

**Figure 4.**
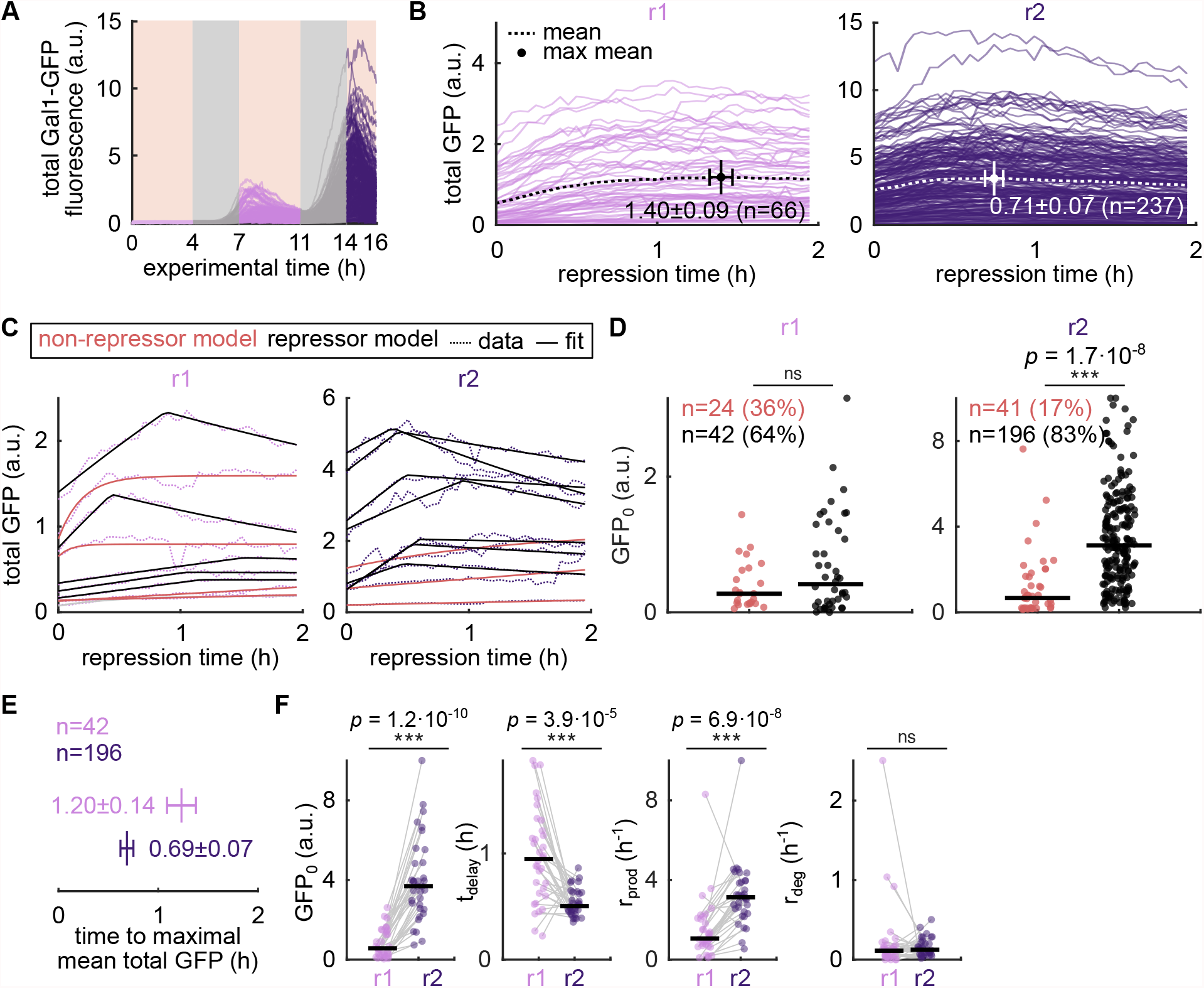
Earlier repression response in repression r2 for *elp6Δ* cells at both the population and single-cell level. (A) Single-cell traces of total Gal1-GFP fluorescence signal of *elp6Δ* budding yeast cells across two inductions i1 and i2 (gray) and repressions 0, 1, and 2 (purple). (B) Single-cell traces of total GFP signal of *elp6Δ* budding yeast cells adjusted for dilution (see Figure 2C) for the first two hours of repression 1 (r1, left) and repression 2 (r2, right). Time to maximal mean total GFP is 41 min shorter in repression r2, where the mean total GFP signal is indicated by the dotted line and the maximal mean total GFP is highlighted by the dot. Bootstrap (10^5^) samples were drawn to generate mean ± std. (C) Ten exemplary total GFP traces (dotted lines) and best fits (solid lines) of *elp6Δ* budding yeast cells and repressions r1 (left) and r2 (right). (D) The median initial total GFP, GFP_0_, is higher in *elp6Δ* traces better fitted by the repressor model (black) than in *elp6Δ* traces better fitted by the non-repressor model (red). This confirms that the repressor model fits induced *elp6Δ* cells better, while the non-repressor model fits uninduced *elp6Δ* cells. The number of cells and percentages of all GFP traces best fitted by the repressor model and non-repressor model are shown in brackets. (E) Time to maximal mean total GFP is decreased in repression r2 for *elp6Δ* repressor cells (1.20 ± 0.14 *vs*. 0.69 ± 0.07). Bootstrap (10^5^) samples of the repressor cells were drawn to generate mean ± std. (F) Comparison of paired estimated single-cell parameters of *elp6Δ* cells of repression r1 and r2 show that median initial total GFP, GFP_0_, median repression delay, t_delay_, and median production rate, r_prod_, are significantly different (*p* = 1.2·10^−10^, *p* = 3.9·10^−5^ and *p* = 6.9·10^−8^, respectively, two-sided paired sign test correcting for multiple testing with Bonferroni correction, m = 20, and the number of paired cells is 34), with both median GFP_0_ and median r_prod_ increased and median t_delay_ decreased (median t_delay_ values of 0.95 and 0.50 h for r1 and r2, respectively) in r2. Median degradation rate, r_deg_, is not significantly different between r1 and r2 (*p* = 0.39). The *elp6Δ* replicate analysis can be found in Figure S2.

### *elp6Δ* shows stronger repression delay impairment in first repression

To first identify if the repression kinetics between wildtype and *elp6Δ* cells differed at the repressor subpopulation level, we compared the repression kinetics of wildtype and *elp6Δ* repressor cells. We found that the time to maximal mean total GFP increased for *elp6Δ* repressor cells for repression r1 (Figure 5A). Next, we wanted to identify if the *elp6Δ* repressor cells show altered repression kinetics at the single-cell level compared to wildtype cells. As we here assess different strains, we were no longer able to pair cells and, thus, compared the estimated single-cell parameters of all repressor cells. Using a two-sided median test and correcting for multiple testing, we discovered that the median t_delay_ for wildtype was significantly lower (median t_delay_ = 0.52h) compared to *elp6Δ* (median t_delay_ = 0.88h, *p* = 4.7·10^−4^) for r1 (Figure 5B). However, median GFP_0_, median r_prod,_ and median r_deg_ are comparable (*p* = 1, *p* = 1, and *p* = 0.44, respectively) (Figure 5B). This is in line with previous findings that wildtype and *elp6Δ* cells have similar Gal1 levels in induction i1 leading to comparable median GFP_0_ in r1 and comparable production rates between wildtype and *elp6Δ* cells during induction i1 (*17*). Then, we performed the same analysis for repression r2. For r2, we found that the time to maximal mean total GFP at the repressor subpopulation level was more similar between wildtype and *elp6Δ* repressor cells (Figure 5C) than in r1. At the single-cell level, the median GFP_0_, median t_delay_, and median r_prod_ were significantly different (*p* = 1.8·10^−6^, *p* = 2.2·10^−3^, and *p* = 8.1·10^−11^, respectively) between wildtype and *elp6Δ* cells for r2 (Figure 5D) (see Materials and Methods), whereas median r_deg_ was comparable after correcting for multiple testing (*p* = 0.04) (see Materials and Methods). Overall, this analysis identifies a - compared to wildtype - stronger repression delay impairment of *elp6Δ* cells in r1, while the repression delay in r2 is more similar to wildtype cells. The previously described gain-of-induction-memory phenotype of *elp6Δ*, which showed identical Gal1 levels to wildtype in induction i1 but increased Gal1 levels in induction i2, is responsible for the increased median GFP_0_ (*17*). Furthermore, Bheda et al. reported increased *elp6Δ* production rates for induction i2 but not for induction i1.

**Figure 5.**
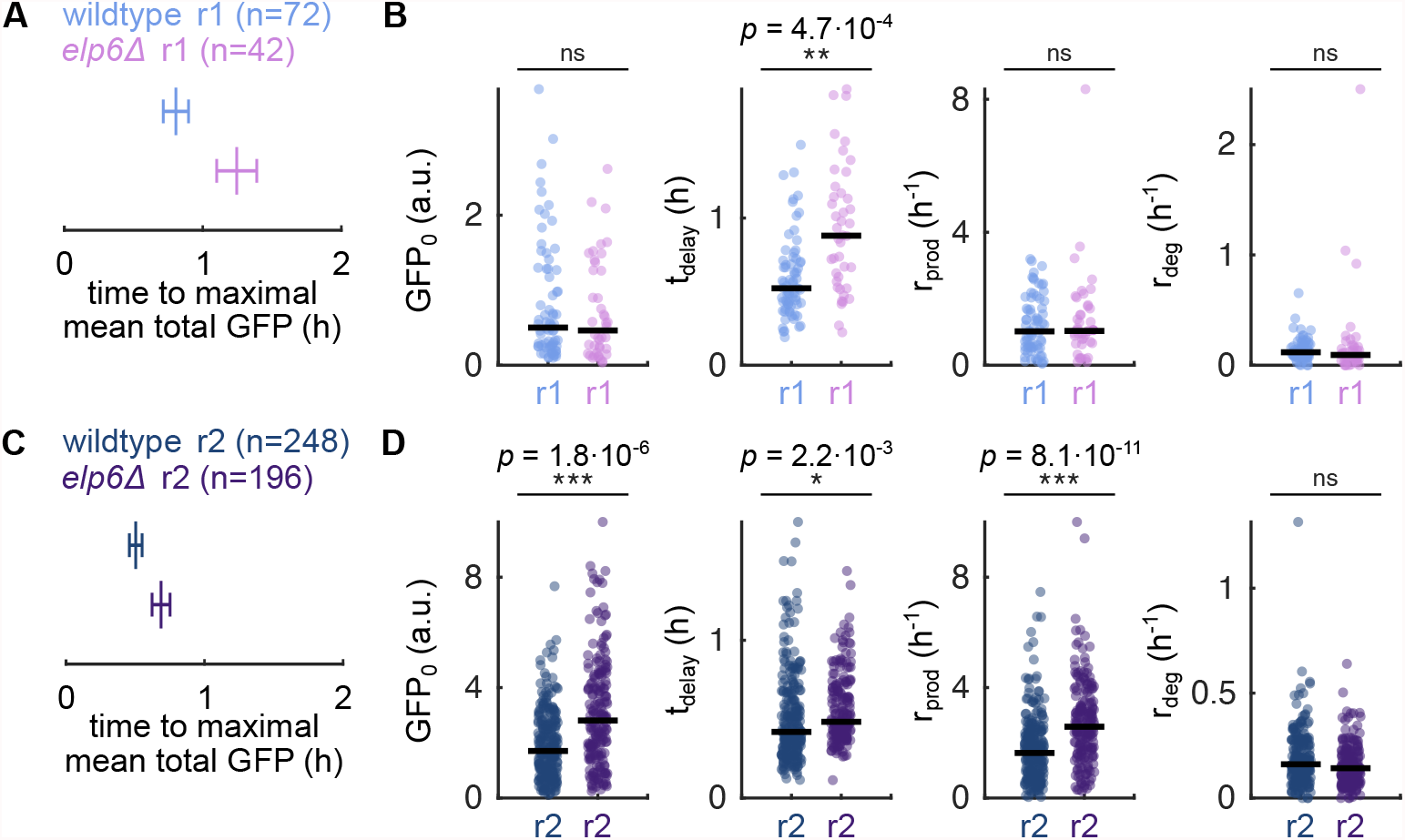
Stronger Gal1 repression delay impairment in *elp6Δ* cells during first repression. (A) Time to maximal mean total GFP is increased for *elp6Δ* repressor cells for repression r1 (0.81 ± 0.09 *vs*. 1.20 ± 0.14). Bootstrap (10^5^) samples were drawn to generate mean ± std. (B) Comparison of estimated single-cell parameters of wildtype and *elp6Δ* cells of repression r1 shows that *elp6Δ* has a significantly different median repression delay, t_delay_, in comparison to wildtype, (*p* = 4.7·10^−4^, two-sided median test correcting for multiple testing with Bonferroni correction, m = 20, and the number of cells for wildtype and *elp6Δ* is 72 and 42, respectively) with median t_delay_ increased in *elp6Δ* (median values of 0.52 and 0.88 h for wildtype and *elp6Δ*, respectively). Median initial total GFP, GFP_0_, median production rate, r_prod_, and median degradation rate, r_deg_, are not significantly different between wildtype and *elp6Δ* cells of repression r1 (*p* =1, *p* = 1, and *p* = 0.44, respectively). (C) Time to maximal mean total GFP is more comparable for wildtype and *elp6Δ* repressor cells for repression r2 (0.52 ± 0.05 *vs*. 0.69 ± 0.07). Bootstrap (10^5^) samples were drawn to generate mean ± std. (D) Comparison of estimated single-cell parameters of wildtype and *elp6Δ* cells of repression r2 shows median GFP_0_, median t_delay_, and median r_prod_, to be significantly different in *elp6Δ* compared to wildtype (*p* = 1.8·10^−6^, *p* = 2.2·10^−3^, and *p* = 8.1·10^−11^, respectively, two-sided median test correcting for multiple testing with Bonferroni correction, m = 20 and number of cells for wildtype and *elp6Δ* is 248 and 196, respectively), with median GFP_0_, median t_delay_, and median r_prod_ increased in *elp6Δ* (median values of 0.42 and 0.48 h for wildtype and *elp6Δ*, respectively). Median r_deg_ is not significantly different between wildtype and *elp6Δ* cells of repression r2 (*p* = 0.04) after correcting for multiple testing.

## DISCUSSION

Based on population data, we found an earlier repression response upon a repeated repression. We used mathematical modeling to demonstrate that changes in the repression response were not simply due to different proportions of repressor cells. Interestingly, we discovered that *elp6Δ* cells showed prolonged repression delay in the first repression but a more comparable delay in comparison to wildtype in the second repression.

### Repression memory manifests as faster initiation of second repression

At both the population and single-cell level, we identified that both wildtype and *elp6Δ* cells have an earlier repression response in repression r2 (Figures 3E–F, 4E–F, S1E-F, S2E-F). This implies that cells repeatedly exposed to glucose are faster at initiating Gal1 repression. We thus propose that budding yeast cells do not only show reinduction memory when exposed to repeated galactose inductions (*17*) but also have a repression memory when exposed to repeated glucose repressions. We hypothesize that faster repression of the galactose-metabolizing machinery would save energy and/or allow for faster induction of the glucose-metabolizing machinery, which would be beneficial to individual yeast cells. It will be intriguing to investigate how Gal1 repression is acquired on a molecular level and if glucose and galactose repression memories are linked in the future. We would like to note here that we assessed the kinetics of the Gal1-GFP fusion protein, which has been successfully employed in several studies involving Gal1 gene expression (*16, 23–26*), and hence also modeled Gal1-GFP kinetics. Moreover, it should be mentioned that potential photo-bleaching effects would not influence the estimated repression delays, and, thus, our repression memory hypothesis. When considering memory as the fold change of the second delay to the first delay, we discover reinduction memory to have an overall stronger effect size. The fold changes of the repression delays are (median t_delay_ r2/median t_delay_ r1 = 0.38/0.50 = 0.76) 0.76 for wildtype and (median t_delay_ r2/median t_delay_ r1 = 0.50/0.95 = 0.53) 0.53 for *elp6Δ*. In comparison, the fold changes of the induction delays are (median t_delay_ i2/median t_delay_ i1 = 0.67/2.33 = 0.29) 0.29 and (median t_delay_ i2/median t_delay_ i1 = 0.50/2.50 = 0.20) 0.20 for wildtype and elp6Δ, respectively, as reported in Bheda et al. (*17*).

### *elp6Δ* is a novel gain-of-repression-memory mutant

We observed that *elp6Δ* cells had longer repression delay in repression r1 than wildtype cells, while the repression delay is more comparable for wildtype and *elp6Δ* cells in repression r2 at both the population and single-cell level (Figure 5). This reveals that the repression memory effects (shortening of repression delay in r2) of *elp6Δ* cells are stronger than in wildtype cells. Hence, *elp6Δ* can be classified as a gain-of-repression-memory mutant. *Elp6* has been identified as one of the six subunits of the so called RNA polymerase II elongator complex (*24*) and has been linked to a variety of biological functions (25–27). However, its precise function is still debated. Contrary to induction, where *elp6Δ* and wildtype show comparable induction delays in the first induction but exhibit significantly different induction delays in the second induction, *elp6Δ* and wildtype cells show significantly different repression delays in r1 and more comparable repression delays in r2 (*17*) (Figure 5B and D). Although reinduction memory has a stronger overall effect size than repression memory, the relative memory effect of *elp6Δ* and wildtype is comparable between induction (*elp6Δ*induction memory/wildtype induction memory = 0.20/0.29 = 0.69) and repression (*elp6Δ* repression memory/wildtype repression memory = 0.53/0.76 = 0.70). Whether the gain-of-repression memory is a unique characteristic of *elp6Δ* cells or if other knock-out mutants show similar repression kinetics and delay responses will be interesting to investigate in the future.

## ACKNOWLEDGMENTS

L.S. was funded by the BMBF project TIDY (031L0170B). L.S. is especially grateful to the Technical University of Munich’s Department of Mathematics, whose Entrepreneurial Award (within the Program “Global Challenges for Women in Math Science”) contributed to the completion of this project. L.S. acknowledges further support by the Add-on Fellowship for Interdisciplinary Life Science of the Joachim Herz Foundation. C.M. acknowledges funding from the European Research Council (ERC) under the European Union’s Horizon 2020 research and innovation program (grant agreement no. 866411). The work in R.S. laboratory was supported by the Deutsche Forschungsgemeinschaft (DFG, German Research Foundation) through SFB 1064 (Project-ID 213249687) and SFB 1309 (Project-ID 325871075), the AmPro program (ZT0026) and Helmholtz Gesellschaft.

## AUTHOR CONTRIBUTIONS

Conceptualization, P.B., R.S. and C.M.; Methodology, L.S.; Software, L.S.; Validation, L.S.; Formal Analysis, L.S.; Investigation, L.S., I.K.; Resources, R.S., and C.M.; Data Curation, L.S., M.B.; Writing – Original Draft, L.S.; Writing – Review & Editing, L.S., M.S., R.S. and C.M.; Visualization, L.S.; Supervision, P.B., R.S., and C.M.; Project Administration, R.S. and C.M.; Funding Acquisition, R.S., and C.M..

## MATERIALS AND METHODS

### Data acquisition and sources

For the analysis for Figures 1-5, we used microscopy images and initial segmentation, mapping, and tracking information from a microfluidics experiment from Bheda et al. (*17*), which contained 13 and ten positions for wildtype and *elp6Δ*, respectively. Positions were chosen according to the feasibility of analysis with respect to cell movement, number and morphology, where each position represents a biological replicate. The images from the first two hours of repression r1 were rectified, and the segmentation, mapping, and tracking were extended to the entire two hours of repression r2. Bheda et al. only segmented r2 partially since they were primarily interested in galactose induction, and did not adjust the final repression frames. Using the software PhyloCell (*19*), we manually corrected the segmentation, mapping, and tracking of r1 and r2 for wildtype and *elp6Δ*. For the technical replicate analysis (Figures S1 and S2) we repeated the induction-repression experiment as described in Bheda et al. (*17*) (Figure 2A). Due to low cell numbers, we pooled data from three and two independent experiments for wildtype and *elp6Δ*, totaling 13 and 23 positions, respectively. Using the software YeaZ (*20*) and Cell-ACDC (*21*) for cell segmentation, mapping and tracking, we extracted the relevant single-cell information of the live-cell images for both repressions r1 and r2. During glucose repression, the yeast cells proliferated, increasing the cell numbers within the microfluidic chambers. However, filled microfluidic chambers no longer assure that all the progeny of a cell is recorded, and mapping and tracking of cells become infeasible. To ensure mapping and tracking of single yeast cells within the microfluidic chambers, the glucose repressions were limited to a maximum of 4 h and the overall experiment was limited to 16 h (4 h in glucose (r0), 3 h in galactose (i1), 4 h in glucose (r1), 3 h in galactose (i2), 2 h in glucose (r2)).

### Data preprocessing

We extracted the single-cell information relevant for our analysis, namely cell ID, mother cell ID, detection frame (first frame in which a cell is detected), last frame (last frame a cell is detected), relative GFP intensities per time (mean GFP intensity of a segmented cell) and cell area per time. As the data regarding the relative GFP intensities and cell area was not sorted over time, we first sorted it and then calculated the total GFP fluorescence per time given by

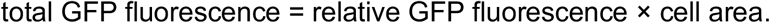

Finally, cells that were not imaged till the end of the experiment, cells with missing relative GFP and/or cell area values, and cells that were supposedly detected before their mother cells (segmentation error) were discarded.

### Dilution compensation

Cytoplasmic proteins are disseminated between the mother and daughter cells during budding. Assuming that Gal1 is not produced or degraded, protein redistribution causes a drop in total Gal1-GFP fluorescence in the mother cell and a rise in total Gal1-GFP fluorescence in the daughter cell till the mother and daughter cells split (Figure 2C top). As a result, regardless of repression, dilution causes variations in total Gal1-GFP fluorescence. The daughter cell grows to about ⅓ of the size of the mother cell (*27*) such that the decrease in total Gal1-GFP fluorescence due to dilution was expected to be ⅓ of the initial total Gal1-GFP fluorescence of the mother cell. To ensure that dilution does not overshadow potentially more subtle repression kinetics, we created artificial non-dividing cells compensating for dilution by adding the total Gal1-GFP fluorescence of the progeny of a cell present at the start of glucose repression, which we called mother cell, to the total Gal1-GFP fluorescence of that mother cell during the first 2 h of repression (Figures 2C bottom, 2D, 4B, S1B, and S2B). For mother cells with a bud at the beginning of a repression period, we additionally added the bud to the total Gal1-GFP fluorescence of that mother cell. The GFP traces of all computed non-dividing cells can be found under https://github.com/marrlab/Gal1repression. As we found the maximal mean total GFP to be attained before 2 h of glucose exposure, we restricted our analysis to the first 2 h of repression.

### Models

During the first two hours of glucose repression, we modeled the kinetics of the total GFP of every single cell. Due to the high variability in galactose induction, we assumed that our initial cell population at the beginning of repression contained induced and uninduced cells, which show and do not show repression kinetics, respectively. We developed two models, the repressor and the non-repressor model, to account for both total GFP kinetics during repression.

#### Non-repressor model

For more information regarding the non-repressor model, see the main text.

#### Repressor model

For more information regarding the repressor model, see the main text.

#### Noise model

Experimental data, such as total GFP per cell per time, is noise corrupted. As a result, we used an underlying additive Gaussian noise model with a constant variance σ^2^ throughout time to test our models. The single-cell specific model parameters are comprised in the parameter vector Θ_i_ for cell i and the experimental measurement at time point k for cell i is denoted by <u>*y*</u>_i_^k^. The log-likelihood for the Gaussian noise model is given by

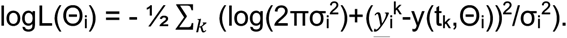

We obtained the optimal model parameters of both models for the total GFP traces for each cell by performing maximum likelihood estimation.

### Optimization and parameter estimation

For each total GFP trace separately for r1 and r2, wildtype, and *elp6Δ*, we computed the model parameters for both the non-repressor and repressor models. The initial total GFP GFP_0_, the basal production rate r_basal_, and the degradation rate r_deg_ are the model parameters for the non-repressor model. Instead of a basal production rate, r_basal_, we have a production rate, r_prod_, for the repressor model. Also, we discovered the time point of delayed repression t_delay_ for the repressor model. For both models we also infer one noise parameter σ determining the spread of the Gaussian noise model. We assumed that all parameters are constant over time. For numerical reasons we optimized the parameters in log_10_ scale (*28*) and rescaled the data by 10^7^. As total Gal1-GFP fluorescence signal and total Gal1-GFP molecules are (linearly) mapped by an unknown constant, the number of total Gal1-GFP molecules is always scalable by that unknown constant that we exploit to increase convergence. The lower and upper bounds for all initial, rate, and noise parameters are –10 and 1 in log_10_ scale, respectively, assuring that the whole range of biologically plausible parameter values is covered. The lower and upper bounds for the repression delay are given by 36 s and 2 h (corresponding to –2 and log_10_(2) in log_10_ scale). As we only considered 2 h of glucose repression, we did not allow the time delay to take on larger values. We performed multi-start local optimization of the negative log-likelihood using the parameter estimation toolbox PESTO (*29*). For each model and total GFP trace, we performed local optimization runs from at least 20 different Latin-hypercube-sampled starts. If less than five starts converged, i.e. the objective function values of the starts differ less than 0.1 to the best start, we re-ran the optimization with 50, 100, and 200 starts until at least five starts converged for each GFP trace.

### Model selection

We used the Bayesian information criterion (BIC) (*30*) for comparing the non-repressor model and the repressor model per total GFP trace. The BIC is calculated by

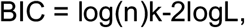

where n is the number of data points, k is the number of estimated parameters and logL is the log-likelihood value for the maximum likelihood estimate of the model parameters. Here, the number of estimated parameters is either four for the non-repressor model or five for the repressor model. The BIC rewards high likelihood values and penalizes the model complexity in the form of additional model parameters. We considered the repressor model to fit a given total GFP trace considerably better than the non-repressor model if BIC_repressor_ < BIC_non-repressor_–10 (Figures 3C–D, 4C–D, S1C-D and S2C-D).

### Statistical analysis

#### Comparison of initial total GFP of total GFP traces

On the estimated initial total GFP, GFP_0_, of all total GFP traces significantly better fitted by a repressor model and all total GFP traces better fitted by a non-repressor model, we did a two-sided median test. To avoid false-positive results, we used the Bonferroni correction, which adjusts the significance-level α = 0.05 by the total number of investigated null hypotheses m, such that α’ = α/m. In this study, the total number of null hypotheses for the main analysis is m = 20:

- Two hypothesis tests comparing initial total GFP between repressor and non-repressor fits for wildtype r1 and r2,
- Four hypothesis tests comparing estimated single-cell parameters between wildtype r1 and r2,
- Two hypothesis tests comparing initial total GFP between repressor and non-repressor fits for *elp6Δ* r1 and r2,
- Four hypothesis tests comparing estimated single-cell parameters between *elp6Δ* r1 and r2,
- Four hypothesis tests comparing estimated single-cell parameters between wildtype and *elp6Δ* for r1, and
- Four hypothesis tests comparing estimated single-cell parameters between wildtype and *elp6Δ* for r2.

In this study, the total number of null hypotheses for the replicate analysis is m = 12:

- Two hypothesis tests comparing initial total GFP between repressor and non-repressor fits for wildtype r1 and r2,
- Four hypothesis tests comparing estimated single-cell parameters between wildtype r1 and r2,
- Two hypothesis tests comparing initial total GFP between repressor and non-repressor fits for *elp6Δ* r1 and r2, and
- Four hypothesis tests comparing estimated single-cell parameters between *elp6Δ* r1 and r2.

#### Comparison of estimated single-cell parameters between repression r1 and r2 for wildtype and elp6Δ

We ignored all traces that were well described by a non-repressor model (uninduced cells) and focused the statistical analysis on the total GFP traces for which the repressor model gave a considerably better fit (see the Model selection, Figures 3C, 4C, S1C and S2C). For those total GFP traces, we compared the estimated single-cell parameters of initial total GFP, GFP_0_, repression delay t_delay_, production and degradation rates, r_prod_ and r_deg_, for repressions r1 and r2 and wildtype and *elp6Δ*. We performed a two-sided paired sign test on the estimated single-cell parameters of paired mother cells in r1 and r2 for both wildtype (Figures 3F and S1F) and *elp6Δ* (Figures 4F and S2F). To avoid false-positive outcomes, we used the Bonferroni correction, which modified the significance level of α = 0.05 by the total number of tested null hypotheses m to α’ = α/m, with m = 20 for the main analysis and m = 12 for the replicate analysis.

#### Comparison of estimated single-cell parameters between wildtype and elp6Δ for repression r1 and r2

We performed a two-sided median test on the estimated single-cell parameters of all cells for wildtype and *elp6Δ* for r1 (Figure 5B) and r2 (Figure 5D) of the main analysis. Due to the comparison between two different yeast strains, we could not perform a paired test. To counteract false-positive results, we again corrected for multiple testing according to the Bonferroni correction, where the significance-level α = 0.05 was adjusted by the total number of tested null hypotheses m to α’ = α/m with m = 20 for the main analysis. As the replicate analysis was performed on pooled data to increase overall cell numbers, we were not able to compare the estimated single-cell parameters between wildtype and *elp6Δ*.

### Implementation

The toolboxes used for segmentation, mapping, and tracking are available at https://github.com/gcharvin/phyloCell (PhyloCell (*19*)), https://github.com/lpbsscientist/YeaZ-GUI (YeaZ (*20*)), and https://github.com/SchmollerLab/Cell_ACDC (Cell-ACDC (*21*)). The toolbox used for parameter estimation (PESTO (*29*)) is available under https://github.com/ICB-DCM. The MATLAB code corresponding to this manuscript is available under https://github.com/marrlab/Gal1repression. The analysis was performed with MATLAB 2017b.

---

**Figures 2 and 3 – Figure S1.**
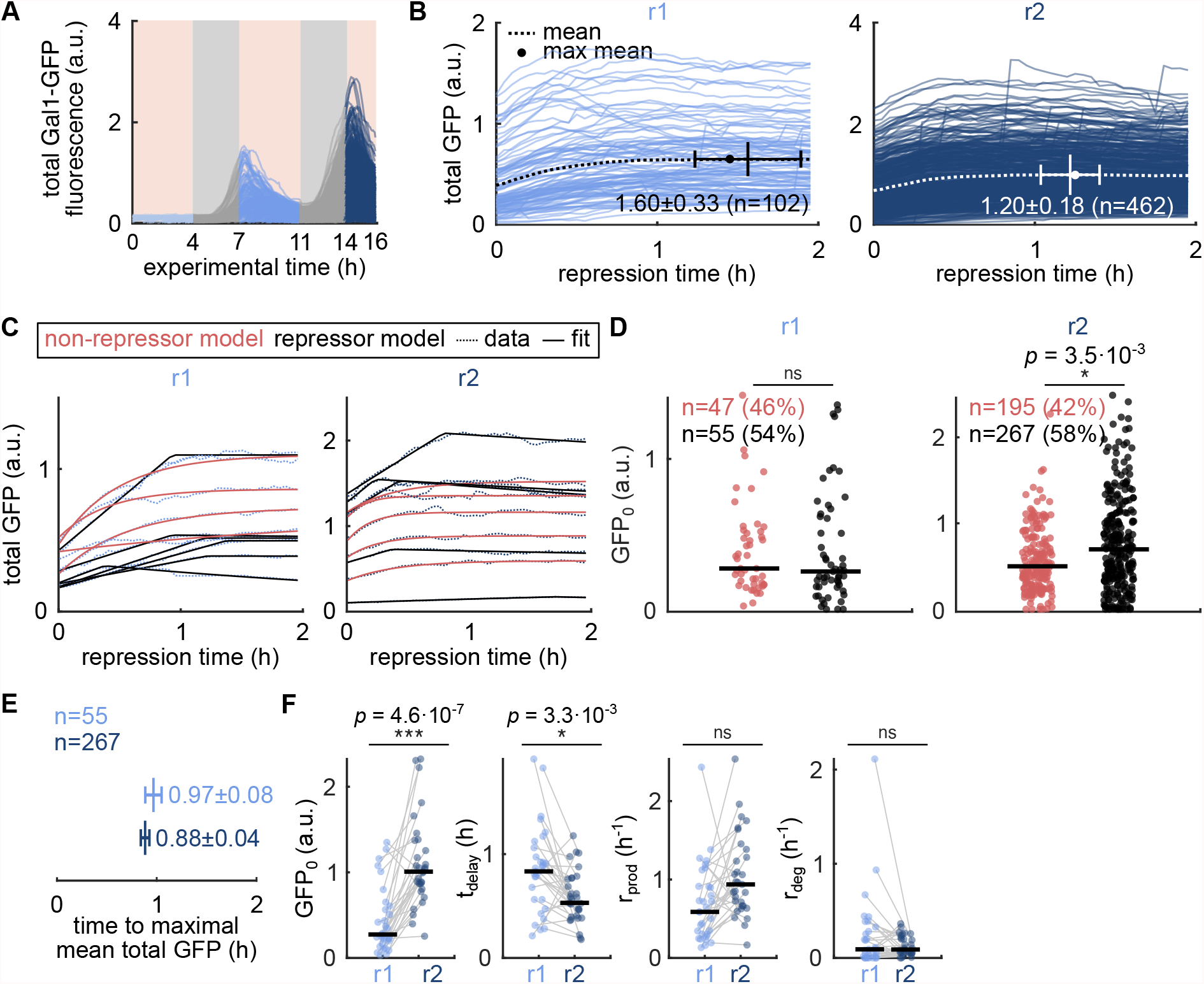
Earlier repression response in repression r2 at both the population and single-cell level for cells of replicate experiment. (A) Single-cell traces of total Gal1-GFP fluorescence signal across two inductions i1 and i2 (gray) and repressions r0, r1, and r2 (blue). (B) Single-cell traces of total GFP signal adjusted for dilution (see (C)) for the first two hours of repression r1 (left) and repression r2 (right). Time to maximal mean total GFP is 24 min shorter in repression r2, where mean total GFP is indicated by the dotted line and the maximal mean total GFP is highlighted by the dot. Bootstrap (10^5^) samples were drawn to generate mean ± std. (C) Ten exemplary total GFP traces (dotted lines) and best fits (solid lines) for repressions r1 (left) and r2 (right). GFP traces best fitted with a repressor model are shown in black and fits of total GFP traces best fitted with a non-repressor model are shown in red. (D) The median initial total GFP, GFP_0_, is higher in traces better fitted by the repressor model (black) than in traces better fitted by the non-repressor model (red). This confirms that the repressor model fits induced cells better, while the non-repressor model fits uninduced cells. The number of cells and percentages of all GFP traces best fitted by the repressor model and non-repressor model are shown. (E) Time to maximal mean total GFP is decreased in repression r2 for repressor cells (0.97 ± 0.08 vs. 0.88 ± 0.04). Bootstrap (10^5^) samples of the repressor cells were drawn to generate mean ± std. (F) Comparison of paired estimated single-cell parameters of repression r1 and r2 shows that the median initial total GFP, GFP_0_, and median repression delay, t_delay_, are significantly different (*p* = 4.6·10^−7^ and *p* = 3.3·10^−3^, respectively, two-sided paired sign test correcting for multiple testing with Bonferroni correction, m = 12, and the number of paired cells = 31), with median GFP_0_ increased and median t_delay_ decreased (median t_delay_ values of 0.83 and 0.53 for r1 and r2, respectively). Median production rate, r_prod_, and median degradation rate, r_deg_, are not significantly different between r1 and r2 (*p* = 0.01 and *p* = 0.50, respectively).

**Figure 4 - Figure S2.**
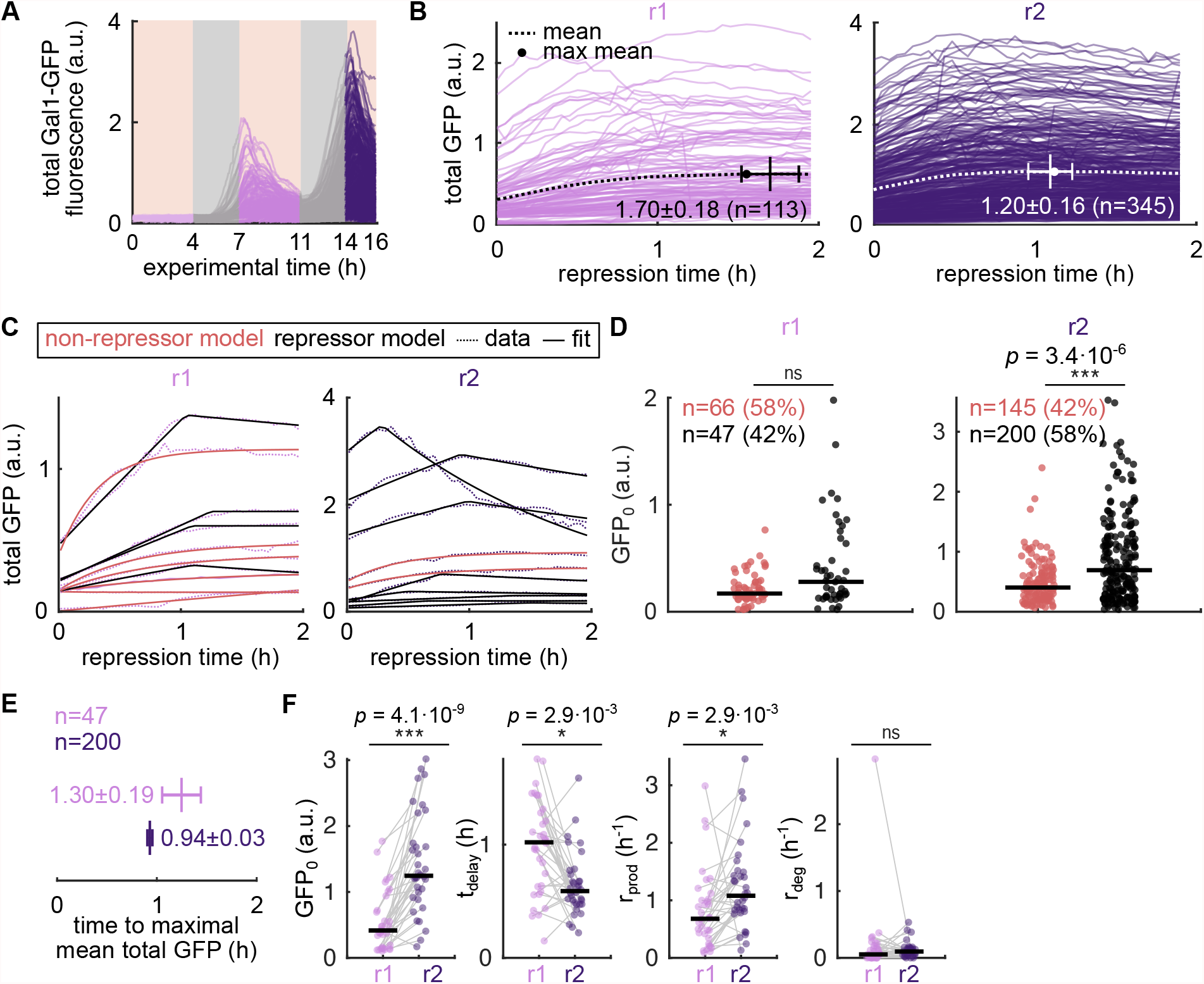
Earlier repression response in repression r2 at both the population and single-cell level for *elp6Δ* cells of replicate experiment. (A) Single-cell traces of total Gal1-GFP fluorescence signal of *elp6Δ* budding yeast cells across two inductions i1 and i2 (gray) and repressions 0, 1, and 2 (purple). (B) Single-cell traces of total GFP signal of *elp6Δ* budding yeast cells adjusted for dilution (see Figure 2C) for the first two hours of r1 (left) and r2 (right). Time to maximal mean total GFP is 36 min shorter in repression r2, where the mean total GFP signal is indicated by the dotted line and the maximal mean total GFP is highlighted by the dot. Bootstrap (10^5^) samples were drawn to generate mean ± std. (C) Ten exemplary total GFP traces (dotted lines) and best fits (solid lines) of *elp6Δ* budding yeast cells and repressions r1 (left) and r2 (right). (D) The median initial total GFP, GFP_0_, is higher in *elp6Δ* traces better fitted by the repressor model (black) than in *elp6Δ* traces better fitted by the non-repressor model (red). This confirms that the repressor model fits induced *elp6Δ* cells better, while the non-repressor model fits uninduced *elp6Δ* cells. The number of cells and percentages of all GFP traces best fitted by the repressor model and non-repressor model are shown. (E) Time to maximal mean total GFP is decreased in repression r2 for *elp6Δ* repressor cells (1.30 ± 0.19 *vs*. 0.94 ± 0.02). Bootstrap (10^5^) samples of the repressor cells were drawn to generate mean ± std. (F) Comparison of paired estimated single-cell parameters of *elp6Δ* cells of repression r1 and r2 show that median initial total GFP, GFP_0_, median repression delay, t_delay_, and median production rate, r_prod_, are significantly different (*p* = 4.1·10^−9^, *p* = 2.9·10^−3^, and *p* = 2.9·10^−3^, respectively, two-sided paired sign test correcting for multiple testing with Bonferroni correction, m = 12, and the number of paired cells is 34), with median GFP_0_ and r_prod_ increased and median t_delay_ decreased (median values of 1.00 and 0.59 h for r1 and r2, respectively) in r2. Median degradation rate, r_deg_, is not significantly different between r1 and r2 (*p* = 0.39).

